# *In silico* design and validation of commercial kit GPS™ CoVID-19 dtec-RT-qPCR Test under criteria of UNE/EN ISO 17025:2005 and ISO/IEC 15189:2012

**DOI:** 10.1101/2020.04.27.065383

**Authors:** Antonio Martínez-Murcia, Gema Bru, Aaron Navarro, Patricia Ros-Tárraga, Adrián García-Sirera, Laura Pérez

## Abstract

**Background:** The Corona Virus Disease 2019 (COVID-19), caused by Severe Acute Respiratory Syndrome Coronavirus 2 (SARS-CoV-2), has become a serious infectious disease affecting human health worldwide and rapidly declared a pandemic by WHO. Early, several RT-qPCR were designed by using only the first SARS-CoV-2 genome sequence.

**Objectives:** A few days later, when additional SARS-CoV-2 genome were retrieved, the kit GPS™ CoVID-19 dtec-RT-qPCR Test was designed to provide a highly specific detection method and commercially available worldwide. The kit was validated following criteria recommended by the UNE/EN ISO 17025:2005 and ISO/IEC 15189:2012.

**Methods:** The present study approached the *in silico* specificity of the GPS™ CoVID-19 dtec-RT-qPCR Test and RT-qPCR designs currently published. The empirical validation parameters specificity (inclusivity/exclusivity), quantitative phase analysis (10-10^6^ copies), reliability (repeatability/reproducibility) and sensitivity (detection/quantification limits) were evaluated for a minimum of 10-15 assays. Diagnostic validation was achieved by two independent reference laboratories, the Instituto de Salud Carlos III (ISCIII), (Madrid, Spain) and the Public Health England (PHE; Colindale, London, UK).

**Results:** The GPS™ RT-qPCR primers and probe showed the highest number of mismatches with the closet related non-SARS-CoV-2 coronavirus, including some indels. The kits passed all parameters of validation with strict acceptance criteria. Results from reference laboratories 100% correlated with these obtained by suing reference methods and received an evaluation with 100% of diagnostic sensitivity and specificity.

**Conclusions:** The GPS™ CoVID-19 dtec-RT-qPCR Test, available with full analytical and diagnostic validation, represents a case of efficient transfer of technology being successfully used since the pandemic was declared. The analysis suggested the GPS™ CoVID-19 dtec-RT-qPCR Test is the more exclusive by far.

## 1. INTRODUCTION

Last 30^th^ January, the Emergency Committee of the World Health Organization (WHO) under the International Health Regulations (IHR) declared an outbreak of pneumonia, lately named Corona Virus Disease 2019 (COVID-19), as a “Public Health Emergency of International Concern” (PHEIC). The disease is caused by Severe Acute Respiratory Syndrome Coronavirus 2 (SARS-CoV-2) and the first genome was rapidly provided (http://virological.org/t/novel-2019-coronavirus-genome/319). SARS-CoV-2 is a Betacoronavirus subgenus *Sarbecovirus* of group 2B and, in many ways, it resembles SARS-CoV, Bat-SARS-CoV and other Bat SARS-like-CoV [1–6] A few weeks later, this novel coronavirus spread worldwide and forced the WHO to declare a Pandemic on March 11 when more than 118,000 positives and 4,291 deaths were already registered in 114 countries. Today, 5^th^ May, the number of positive cases globally amounts to more than 3.6 million people with more than 250,000 deaths. Faced with the aggressiveness of this global alarm, the massive, reliable, and rapid diagnosis is undoubtedly vital and foremost priority for decision-making at each stage to facilitate public health interventions, and the needs have overwhelmed any forecast.

Current molecular diagnostic tools for viral detection are typically based upon the amplification of target-specific genetic sequences using the Polymerase Chain Reaction (PCR). In acute respiratory infection, real time PCR (so-called quantitative PCR; qPCR) is the gold-standard and routinely used to detect causative viruses as, by far, is the most sensitive and reliable method [7–11]. On the 17^th^ January, the WHO published the very first primers and probes for Reverse Transcriptase qPCR (RT-qPCR) developed by Corman et al., 2020 [12]. They used known genomic data from SARS-CoV and SARS-CoV related (Bat viruses) to generate a non-redundant alignment. The candidate diagnostic RT-PCR assay was designed upon the first SARS-CoV-2 sequence release, based on the sequence alignment match to known SARS-CoV. Because only a single SARS-CoV-2 genome was available, the two monoplex PCR protocols (ORF1ab and N genes) designed to detect SARS-CoV-2 are also reactive to SARS-CoV and Bat SARS-like-CoV. A few days later, 23^rd^ January, the same laboratory together with reference laboratories from the Netherlands, Hong Kong, France, United Kingdom, and Belgium, added a third monoplex-RT-qPCR [12]. Many laboratories worldwide are currently using this RT-qPCR protocol [13] and also it has been the basis to develop many commercial kits. Almost simultaneously, other primers and probes were designed and available by scientists from the Institut Pasteur, París; Centers for Disease Control and Prevention (CDC), Division of Viral Diseases, Atlanta, USA; National Institute for Viral Disease Control and Prevention (CDC), China; Hong Kong University; Department of Medical Sciences, Ministry of Public Health, Thailand; the National Institute of Infectious Diseases, Japan [12–19]. The Respiratory Viruses Branch, Division of Viral Diseases, CDC, Atlanta, recently (4^th^ February) updated a manual of Real-Time RT-PCR Panel for detection of this 2019-Novel Coronavirus (SARS-CoV-2), which was modified 15^th^ March. The SARS-CoV-2 primer and probe sets were designed for the universal detection of SARS-like coronaviruses (N3 assay) and for specific detection of SARS-CoV-2 (N1 and N2 assays). Finally, authors from the Institut Pasteur, Paris, based on the first sequences of SARS-CoV-2 available on the GISAID database (Global Initiative on Sharing All Influenza Data) on 11^th^ January, updated a protocol for the detection of SARS-CoV-2 for two RdRp targets (IP2 and IP4) [14].

Some biotechnology-based companies have recently developed kits for detection of SARS-CoV-2, based on RT-qPCR and provided easy transfer of technology to laboratories worldwide. A fully SARS-CoV-2-specific RT-qPCR thermostable kit was early launched on 27^th^ January by Genetic PCR Solutions™ (GPS™), a brand of Genetic Analysis Strategies SL. (Alicante, Spain). The alignments used at that time included 13 SARS-CoV-2 genome sequences released by 6 different laboratories, deposited in GISAID and available since 19^th^ January 2020. With the purpose to discriminate this new SARS-CoV-2 of present outbreak from previous related SARS, a second independent monoplex RT-qPCR test to detect any other non-SARS-CoV-2 was also produced and provided (not shown). On this study, we have performed a deep analytical and diagnostic validation of the GPS™ COVID-19 dtec-RT-qPCR Test, following the UNE/EN ISO 17025:2005 and ISO/IEC 15189:2012, respectively. A comparative analysis of the specificity (inclusivity and exclusivity) of the designed primers and probes with most previously published RT-qPCR methods is also here reported.

## 2. MATERIALS AND METHODS

### 2.1. GENOME SEQUENCES ALIGNMENT AND PHYLOGENETIC ANALYSIS

Partial alignments of ten SARS-CoV-2 genomic sequences and these from strains of Bat-CoV, Bat SARS-like-CoV, SARS-CoV, Pangolin-CoV (ca. 18,141 bp) and the corresponding phylogenetic tree (Figure 1) was obtained by Neighbour joining method [20], with bootstrap values for 1000 replicates, using the MEGA 5.2.2 software [21].

**Figure 1.**
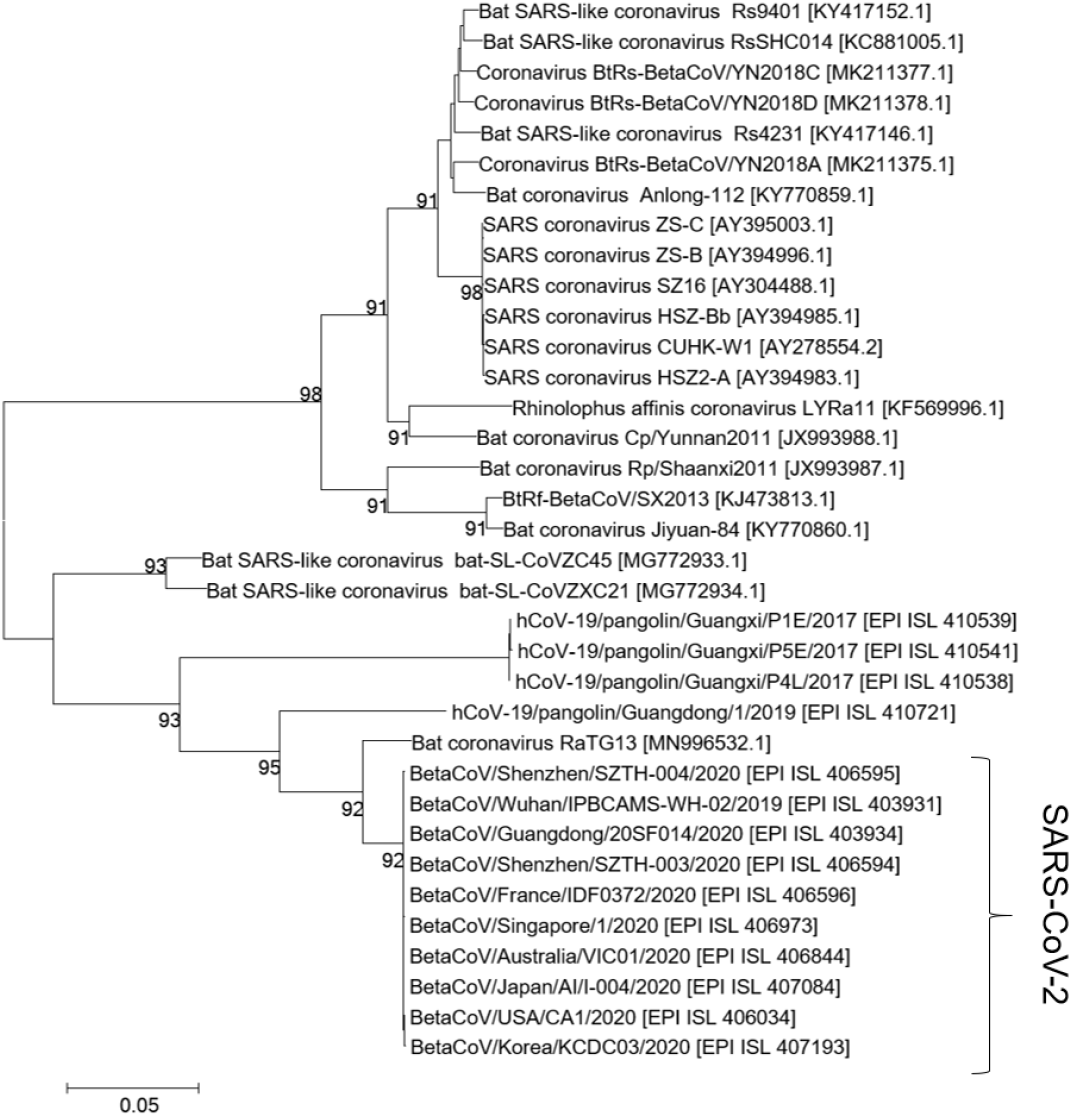
Phylogenetic neighbour-joining tree showing relationships of SARS-CoV-2 and the most related strains of some Betacoronavirus, including SARS-CoV, Bat-CoV, Bat SARS-like-CoV and Pangolin-CoV. The analysis was derived from the alignment of 18,141 nucleotides. Numbers at nodes indicate bootstrap values (percentage of 1000 replicates).

### 2.2. *IN SILICO* COMPARATIVE ANALYSIS OF PRIMERS/PROBES SPECIFICITY

The primers and probes of GPS™ COVID-19 dtec-RT-qPCR Test and the RT-qPCR designs recently published [12–19] were aligned to the corresponding homologous regions of 63 SARS-CoV-2 strains and closely related Betacoronavirus using the Basic Local Alignment Search Tool (BLAST) software available on the National Center for Biotechnology Information (NCBI, https://blast.ncbi.nlm.nih.gov/Blast.cgi) website databases (Bethesda, MD, USA). This *in silico* analysis was periodically updated with new entries currently available. Number of mismatches of the primers and probes of the GPS™ kit and the designs recently published was calculated to evaluate the in silico specificity (Table 1). An illustration of the mismatching of primers/probe sequences of the GPS™ CoVID-19 dtec-RT-qPCR Test, respect of the SARS-CoV-2, Bat SARS-like-CoV, SARS-CoV, Bat-CoV, and Pangolin-CoV groups is shown in Figure 2.

**Figure 2.**
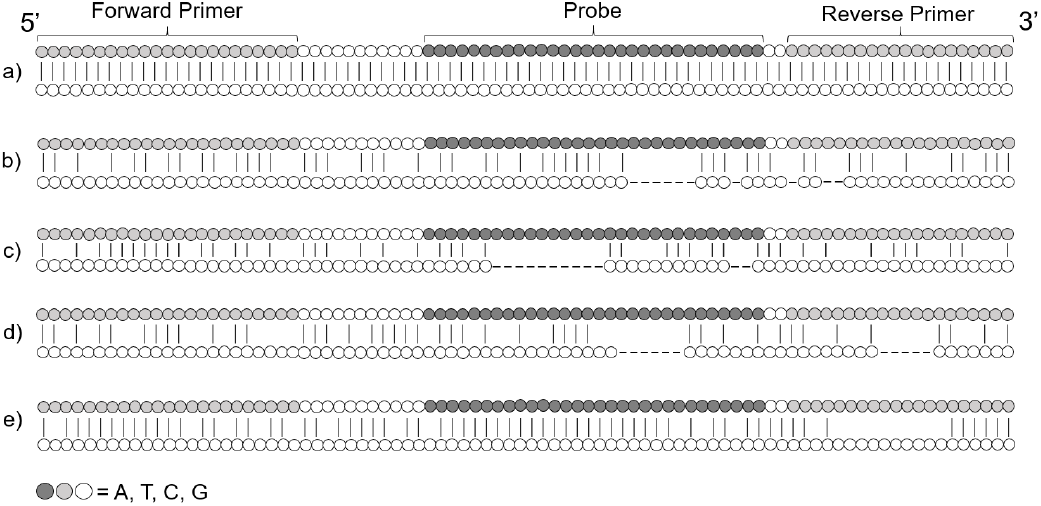
Illustrative alignment representation of the primers/probes sequences of GPS™ CoVID-19 dtec-RT-qPCR Test with a) SARS-CoV-2 (MN975262.1); b) Bat SARS-like-CoV (MG772934.1); c) SARS-CoV (AY304489.1); d) Bat-CoV (KY770859.1); and e) Pangolin-CoV (EPI_ISL_410539).

**Table 1.** Number of mismatches found in the primers/probes sets of the GPS™ COVID-19 dtec-RT-qPCR Test and recently published, from the comparative *in silico* analysis with Bat-CoV, Bat SARS-like-CoV, SARS-CoV and Pangolin-CoV. Numbers in bold show the sum of the mismatches found in the primers/probes of the RT-qPCR designs. Numbers in brackets show the mismatches found in forward primer (FP), probe (P) and reverse primer (RP) following this format: [FP / P / RP].

| Reference | INSTITUTION | TARGET | TOTAL MISMATCHES |  |  |  |  |  |  |  |  |  |
| --- | --- | --- | --- | --- | --- | --- | --- | --- | --- | --- | --- | --- |
|  |  |  | SARS-CoV-2 |  | Bat coronavirus |  | Bat SARS-like coronavirus |  | SARS coronavirus |  | Pangolin coronavirus |  |
|  | Genetic PCR solutions™ (Spain) | - | **0-1** | [0 / 0-1 / 0] | **37-48** | [8-11 / 17-23 / 12-14] | **26-37** | [6-9 / 12-17 / 8-11] | **36-38** | [7-8 / 19 / 9] | **19-31** | [5-9 / 4-11 / 10-11] |
| [14] | Institut Pasteur (Paris) | RdRp (IP2) | **0** | [0 / 0 / 0] | **6-9** | [1-2 / 2-3 / 3-4] | **10-11** | [4 / 2-3 / 4] | **7-12** | [1-4 / 3 / 3-5] | **4-8** | [0-4 / 2 / 2] |
|  |  | RdRp (IP4) | **0** | [0 / 0 / 0] | **12-13** | [1 / 6 / 5-6] | **12-17** | [1-2 / 5-7 / 6-8] | **14** | [2 / 6 / 6] | **4-11** | [0-1 / 2-3 / 2-7] |
| [15] | Centers for Disease Control and Prevention (Atlanta) | N (1) | **0-1** | [0-1 / 0 / 0] | **8-10** | [3-4 / 2 / 3-4] | **8-10** | [3-4 / 2 / 3-4] | **11** | [7 / 2 / 2] | **2-11** | [1-7 / 0-1 / 1-3] |
|  |  | N (2) | **0** | [0 / 0 / 0] | **6** | [0 / 5 / 1] | **6** | [0 / 5 / 1] | **7** | [0 / 5 / 2] | **4** | [1 / 3 / 0] |
|  |  | N (3) | **0-1** | [0 / 0-1 / 0] | **3-5** | [1-2 / 1 / 1-2] | **2-5** | [1-3 / 1 / 0-1] | **4** | [1 / 1 / 2] | **1-3** | [0-1 / 1-2 / 0] |
| [12] | Charité (Berlin) | RdRp_P1 | **3** | [0 / 2 / 1] | **2** | [0 / 2 / 0] | **2-3** | [0-1 / 1 / 1] | **1** | [0 / 1 / 0] | **2-3** | [0-1 / 1 / 1] |
|  |  | RdRp_P2 | **1** | [0 / 0 / 1] | **4** | [0 / 4 / 0] | **3-5** | [0-1 / 2-3 / 1] | **2-3** | [0 / 2-3 / 0] | **2-5** | [0-1 / 1-3 / 1] |
|  |  | E | **0** | [0 / 0 / 0] | **0-3** | [0-1 / 0-1 / 0-1] | **0** | [0 / 0 / 0] | **0-5** | [0-3 / 0-1 / 0-1] | **0** | [0 / 0 / 0] |
| [16] | National Institute for Viral Disease Control and Prevention (China) | ORF1ab | **0** | [0 / 0 / 0] | **7-8** | [2 / 1 / 4-5] | **7-9** | [1-2 / 1-2 / 5] | **7-8** | [2 / 1 / 4-5] | **2-4** | [0-1 / 0-1 / 2] |
|  |  | N | **0** | [0 / 0 / 0] | **8-10** | [2 / 4-5 / 2-3] | **5-10** | [0-3 / 3-4 / 2-3] | **8** | [2 / 4 / 2] | **3-8** | [1 / 1-4 / 1-3] |
| [17] | Hong Kong University Faculty of Medicine (Hong Kong) | ORF1b | **0** | [0 / 0 / 0] | **0-1** | [0 / 0-1 / 0] | **0-1** | [0 / 0-1 / 0] | **0-6** | [0 / 0-2 / 0-4] | **2** | [1 / 0 / 1] |
|  |  | N | **4** | [0 / 4 / 0] | **4** | [0 / 4 / 0] | **4** | [0 / 4 / 0] | **5** | [1 / 4 / 0] | **4-5** | [0-1 / 4 / 0] |
| [18] | Ministry of Public Health (Thailand) | N | **0** | [0 / 0 / 0] | **6** | [1 / 2 / 3] | **6-7** | [1-2 / 2 / 3] | **6** | [2 / 2 / 2] | **2-6** | [0-1 / 1-3 / 1-2] |
| [19] | National Institute of Infectious Diseases (Japan) | N | **1** | [0 / 0 / 1] | **9-12** | [2-4 / 1 / 6-7] | **10** | [2 / 1 / 7] | **11** | [3 / 1 / 7] | **3-7** | [2-4 / 0 / 1-3] |
| [13] | Hong Kong University State Key Laboratory of Emerging Infectious Diseases (Hong Kong) | RdRp/Hel | **1** | [0 / 0 / 1] | **12-18** | [1-2 / 8-12 / 3-4] | **11-15** | [2 / 8-9 / 1-4] | **11** | [1 / 10 / 0] | **4-6** | [1 / 2-3 / 1-2] |
|  |  | S | **0** | [0 / 0 / 0] | **14-22** | [6-8 / 6-9 / 2-5] | **24-25** | [8-9 / 9 / 7] | **23** | [8 / 7 / 8] | **11-19** | [3-7 / 4-8 / 4] |
|  |  | N | **0-1** | [0 / 0 / 0-1] | **10-13** | [2-3 / 7 / 1-3] | **10-11** | [1-2 / 7 / 2] | **11-12** | [2-3 / 7 / 2] | **2-7** | [1 / 0-3 / 1-3] |

### 2.3. GPS™ COVID-19 dtec-RT-qPCR Test

Assays using the GPS™ COVID-19 dtec-RT-qPCR kit (Alicante, Spain) were prepared and reaction mixtures were subjected to qPCR in a QuantStudio3 (ABI) as described in the manual provided. Internal, positive, and negative PCR controls were included. Standard curve calibration of the qPCR was performed by preparing ten-fold dilution series containing 10 to 10^6^ copies of standard template provide in the kit, but also using 5·10^6^ to 5·10 copies of two complete synthetic RNA genomes from SARS-CoV-2 isolate Australia/VIC01/2020 (GenBank No.: MT007544.1) and isolate Wuhan-Hu-1, (GenBank No.: MN908947.3), provided by Twist Bioscience (South San Francisco, United States of America).

### 2.4. ANALYTICAL AND DIAGNOSTIC VALIDATION OF THE GPS™ CoVID-19 dtec-RT-qPCR Test

The method for SARS-CoV-2 detection using the GPS™ kit was subjected to strict validation according to guidelines of the UNE/EN ISO/IEC 17025:2005 and ISO/IEC 15189 [22, 23], as previously described in detail [22]. Validation terms included were repeated 10-15 times *and* the acceptance criteria are shown in Table 2. Diagnostic validation was a service performed by the Instituto de Salud Carlos III (ISCIII), reference laboratory for biomedical investigation and Public Health (Madrid, Spain), by testing 80 breath specimens of the anonymous biobank of Centro Nacional de Microbiología (CNM, Madrid, Spain) previously characterized by a reference protocol [12]. The GPS™ kit was also evaluated by the Public Health England (PHE; Colindale, London, UK) with a sample-panel of 195 specimens, including respiratory clinical specimens negative for SARS-CoV-2 as determined by the validated in-house PHE PCR assay (RdRP gene) and three dilutions of SARS-CoV-2 positive material.

**Table 2.** Summarized results of CoVID-19 dtec-RT-qPCR Test validation according with the guidelines of the UNE/EN ISO/IEC 17025:2005 and ISO/IEC 15189:2012, and acceptance criteria adopted.

| Term of validation | Obtained values |  | Acceptance criteria | Result |
| --- | --- | --- | --- | --- |
| **Specificity** | **Positive:** SARS-CoV-2 isolate Australia/VIC01/2020 (GenBank No.: MT007544.1) and isolate Wuhan-Hu-1 (GenBank No.: MN908947.3) |  | **Inclusiveness:**<br>Positive for both SARS-CoV-2 strains | **ACCEPTED** |
|  | **Negative:** 39 negative specimens from ISCIII, previously characterized by reference protocol [12] |  | **Exclusiveness:**<br>Negative for all negative specimens, | **ACCEPTED** |
| **Standard curve** | $Y = -3.534 \cdot m + 37.534$<br>$a = -3.534$<br>$R^2 = 0.9986$ | | $-3.587 < a < -3.103$ | **ACCEPTED** |
| | $F_{\text{assay}} = 0.014$<br>$F_{\text{fisher}} = 5.318$ | | $F_{\text{assay}} < F_{\text{fisher}}$ | **ACCEPTED** |
| | Efficiency (e) = 93.1 % | | $90 \% < e < 110\%$ | **VALIDATED** |
| **Reliability** | **Repeatability** | | $CV < 10\%$ | **REPEATABLE** |
|  | Conc. | CV (%) |  |  |
| | $10^6$ copies | 1.18 | | |
| | $10^5$ copies | 1.08 | | |
| | $10^4$ copies | 0.68 | | |
| | $10^3$ copies | 0.53 | | |
| | $10^2$ copies | 0.54 | | |
|  | 10 copies | 1.31 |  |  |
| | **Reproducibility** | | $CV < 10\%$ | **REPRODUCIBLE** |
|  | Conc. | CV (%) |  |  |
| | $10^6$ copies | 1.13 | | |
| | $10^5$ copies | 0.91 | | |
| | $10^4$ copies | 0.93 | | |
| | $10^3$ copies | 0.59 | | |
| | $10^2$ copies | 0.66 | | |
|  | 10 copies | 1.83 |  |  |
| **Limit of Detection (LOD)\*** | 10 copies | Positive = 15/15 (100%) | Positives $\geq 90 \%$ | **ACCEPTED** |
| **Limit of Quantification (LOQ)\*** | 10 copies | $t_{\text{value}} = 0.582$<br>$t_{\text{student}} = 2.145$ | $t_{\text{value}} < t_{\text{student}}$ | **ACCEPTED** |
| **Diagnostic Validation** | Diagnostic Specificity: 100%<br>Diagnostic Sensitivity: 100%<br>Diagnostic Efficiency: 100% | | $\geq 90 \%$ | **ACCEPTED** |

## 3. RESULTS

The phylogenetic relationships of selected SARS-CoV-2 genomes and other Betacoronavirus SARS-CoV, Bat SARS-like-CoV, Bat-CoV, and Pangolin-CoV are shown in Figure 1. The analysis indicated that Bat-CoV RaTG13 and a sequence of Pangolin-CoV showed the highest sequence similarity to SARS-CoV-2 (96.70% and 90.74%, respectively) while other Pangolin-CoV sequences available showed a lower homology (85.21%). For *in silico* specificity analysis, primers and probes sequences of the GPS™ kit and the other RT-qPCR designs recently published (16-27), were aligned to SARS-CoV-2 and the other Betacoronaviruses sequences and number of mismatches were annotated in Table 1. In order to illustrate the extent of mismatching of GPS™ kit, an alignment of primers/probe sequences to selected SARS-CoV sequences is shown in Figure 2. Analytical and diagnostic validation of the GPS™ kit, according to the guidelines of the UNE/EN ISO/IEC 17025:2005 and ISO/IEC 15189 [23, 24], was undertaken and the results were summarized in Table 2. Standard calibration curves of the qPCR were performed from ten-fold dilution series (Figure 3a, b) and synthetic RNA genomes of Australia/VIC01/2020 and Wuhan-Hu-1 SARS-CoV-2 isolates (Figure 3c, d). Finally, results of diagnostic validation achieved by the Instituto de Salud Carlos III (ISCIII) are shown in Table 3, and 100% of diagnostic sensitivity and specificity was assigned. Evaluated by the Public Health England (PHE; Colindale, London, UK) and yielded 100% correlation with reference RT-qPCR (not shown).

**Figure 3.**
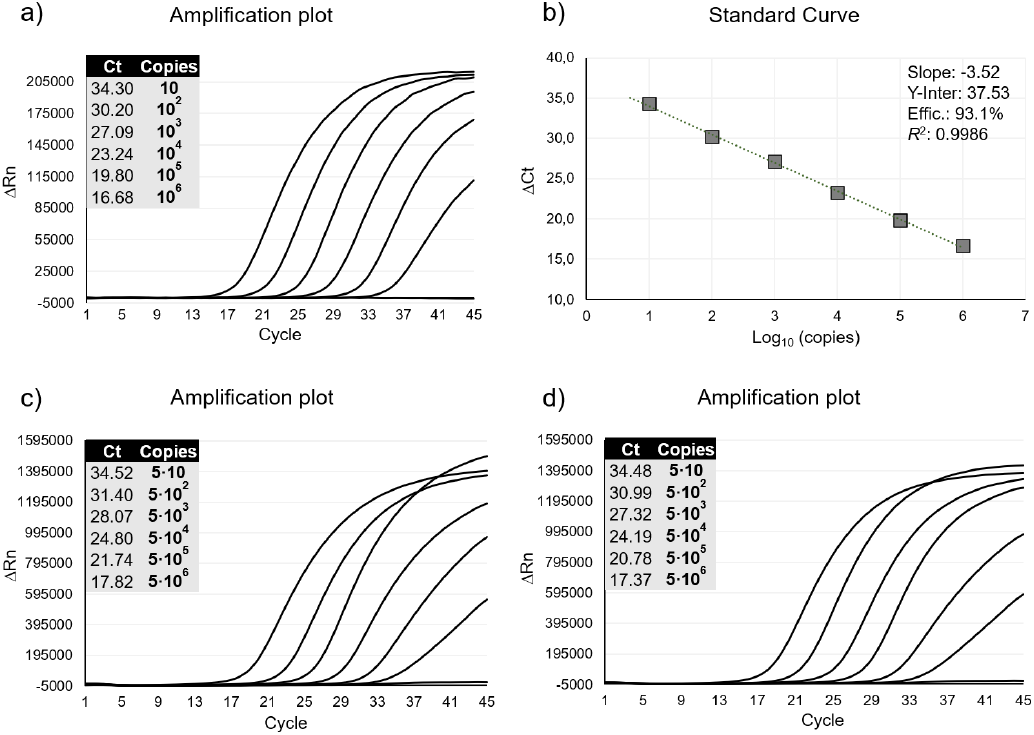
Quality Control of the GPS™ CoVID-19 dtec-RT-qPCR Test with data of six ranges of decimal dilution from 10^6^ copies to 10 copies, and negative control. a) Amplification plot and b) a representative calibration curve with stats. Inclusivity of the GPS™ CoVID-19 dtec-RT-qPCR Test using six ranges of decimal dilution from 5·10^6^ copies to 5·10 copies, and negative control. Amplification plot of synthetic RNA of c) Australian strain of SARS-CoV-2 (MT007544.1); and d) Wuhan-Hu-1 strain of SARS-CoV-2 (MN908947.3).

**Table 3.** Results obtained with GPS™ CoVID-19 dtec-RT-qPCR Test in 80 breath specimens compared with the Ct values determined by using a reference protocol [12], at the Instituto de Salud Carlos III (Madrid)

| CNM Code | CNM Result | CNM Ct GEN1 | CNM Ct GEN2 | CoVID-19 dtec-RT-qPCRT Test | Ct CoVID-19 dtec-RT-qPCR Test |
| --- | --- | --- | --- | --- | --- |
| #01 | NEG | 0 | 0 | NEG | 0 |
| #02 | NEG | 0 | 0 | NEG | 0 |
| #03 | POS | 24 | 28 | POS | 29.43 |
| #04 | POS | 24 | 28 | POS | 23.06 |
| #05 | POS | 23.19 | 26.10 | POS | 27.56 |
| #06 | NEG | 0 | 0 | NEG | 0 |
| #07 | NEG | 0 | 0 | NEG | 0 |
| #08 | POS | 27.16 | 30.41 | POS | 32.18 |
| #09 | NEG | 0 | 0 | NEG | 0 |
| #10 | POS | 20.19 | 25.17 | POS | 21.42 |
| #11 | POS | 28 | 32 | POS | 30.63 |
| #12 | NEG | 0 | 0 | NEG | 0 |
| #13 | POS | 28 | 31 | POS | 30.06 |
| #14 | NEG | 0 | 0 | NEG | 0 |
| #15 | POS | 23.19 | 26.1 | POS | 24.72 |
| #16 | NEG | 0 | 0 | NEG | 0 |
| #17 | POS | 27.48 | 31.16 | POS | 19.15 |
| #18 | NEG | 0 | 0 | NEG | 0 |
| #19 | NEG | 0 | 0 | NEG | 0 |
| #20 | POS | 23 | 25 | POS | 16.07 |
| #21 | POS | 23 | 25 | POS | 19.32 |
| #22 | NEG | 0 | 0 | NEG | 0 |
| #23 | NEG | 0 | 0 | NEG | 0 |
| #24 | POS | 25 | 29 | POS | 24.37 |
| #25 | POS | 20 | 22 | POS | 26.47 |
| #26 | NEG | 0 | 0 | NEG | 0 |
| #27 | POS | 23 | 25 | POS | 24.86 |
| #28 | NEG | 0 | 0 | NEG | 0 |
| #29 | NEG | 0 | 0 | NEG | 0 |
| #30 | POS | 24 | 27 | POS | 23.45 |
| #31 | NEG | 0 | 0 | NEG | 0 |
| #32 | POS | 24.29 | 27.08 | POS | 16.2 |
| #33 | NEG | 0 | 0 | NEG | 0 |
| #34 | NEG | 0 | 0 | NEG | 0 |
| #35 | POS | 27.16 | 30.41 | POS | 29.22 |
| #36 | NEG | 0 | 0 | NEG | 0 |
| #37 | NEG | 0 | 0 | NEG | 0 |
| #38 | POS | 31 | 34 | POS | 33.41 |
| #39 | NEG | 0 | 0 | NEG | 0 |
| #40 | POS | 23.13 | 26.49 | POS | 25 |
| #41 | POS | 16.61 | 19.06 | POS | 16.58 |
| #42 | NEG | 0 | 0 | NEG | 0 |
| #43 | POS | 22.14 | 25.35 | POS | 24.01 |
| #44 | POS | 26.47 | 29.47 | POS | 27.17 |
| #45 | NEG | 0 | 0 | NEG | 0 |
| #46 | POS | 25.59 | 28.03 | POS | 27.23 |
| #47 | POS | 24.16 | 26.44 | POS | 25.96 |
| #48 | NEG | 0 | 0 | NEG | 0 |
| #49 | POS | 24.27 | 26.48 | POS | 25.99 |
| #50 | NEG | 0 | 0 | NEG | 0 |
| #51 | NEG | 0 | 0 | NEG | 0 |
| #52 | NEG | 0 | 0 | NEG | 0 |
| #53 | POS | 24.40 | 26.61 | POS | 26.37 |
| #54 | NEG | 0 | 0 | NEG | 0 |
| #55 | POS | 25.33 | 26.75 | POS | 25.66 |
| #56 | NEG | 0 | 0 | NEG | 0 |
| #57 | NEG | 0 | 0 | NEG | 0 |
| #58 | POS | 25.69 | 28.39 | POS | 27.08 |
| #59 | NEG | 0 | 0 | NEG | 0 |
| #60 | NEG | 0 | 0 | NEG | 0 |
| #61 | POS | 25.73 | 28.61 | POS | 27.24 |
| #62 | POS | 25.91 | 28.43 | POS | 27.46 |
| #63 | NEG | 0 | 0 | NEG | 0 |
| #64 | POS | 26.11 | 28.2 | POS | 27.98 |
| #65 | POS | 25.29 | 28.17 | POS | 27.84 |
| #66 | NEG | 0 | 0 | NEG | 0 |
| #67 | POS | 24.24 | 27.33 | POS | 26.19 |
| #68 | NEG | 0 | 0 | NEG | 0 |
| #69 | NEG | 0 | 0 | NEG | 0 |
| #70 | POS | 24.25 | 26.87 | POS | 26.81 |
| #71 | POS | 26.5 | 29.08 | POS | 26.35 |
| #72 | POS | 25.29 | 28.17 | POS | 26.91 |
| #73 | POS | 26.11 | 28.20 | POS | 26.45 |
| #74 | NEG | 0 | 0 | NEG | 0 |
| #75 | POS | 24.24 | 27.33 | POS | 26.41 |
| #76 | POS | 24.19 | 26.67 | POS | 26.65 |
| #77 | NEG | 0 | 0 | NEG | 0 |
| #78 | POS | 25.23 | 30.18 | POS | 31.72 |
| #79 | NEG | 0 | 0 | NEG | 0 |
| #80 | POS | 24.03 | 26.88 | POS | 26.43 |

## 4. DISCUSSION

Only three months ago, an outbreak of severe pneumonia caused by the novel coronavirus SARS-CoV-2 started in Wuhan (China) and rapidly expanded to almost all areas worldwide. Due to the need of urgent detection tools, several laboratories developed RT-qPCR methods by designing primers and probes from the alignment of a single-first provided SARS-CoV-2 genome sequence to known SARS-CoV, and some of these protocols were published at the WHO website [12–19]. As the number of genomes available rapidly expanded during last January, the GPS™ CoVID-19 dtec-RT-qPCR Test was based on a more specific target for SARS-CoV-2 detection, being this company one of the pioneers marketing a PCR-kit for the CoVID-19 worldwide.

The phylogenetic analysis indicated that, while SARS-CoV-2 shows a high sequence homology (over 99.91-99.97%), the closest relatives were strains of several Betacoronaviruses, with considerable sequence identity to Pangolin isolates, (Figure 1) which confirmed previous results [1, 6, 25–27, 29]. We have found that a single genome sequence of the Bat coronavirus RaTG13 isolated from *Rhinolophus affinis* in Wuhan, showed the highest homology level (96.70%) to SARS-CoV-2, as previously described [1, 2, 4, 6, 29, 2, 30]. However, because only a single Bat-CoV sequence showing this high identity is available, and it was deposited after the outbreak started (27^th^ January), the possibility of RNA contamination during genome sequencing should be ruled-out before take further conclusions. During the design of the GPS™ kit, a purpose of present study was the comparison *in silico* (Table 1) with designed primers and probes so far published [12–19]. In overall, all qPCR designs were inclusive for SARS-CoV-2 as primers and probes showed a good matching. Only the probe for N gene designed by Chu et al., 2020 [17] showed 4 mismatches which may affect to its binding, particularly considering its short primary structure. In some cases, single nucleotide mismatching was observed in some primers, but none of them were located close to primer 3’-end. Considering all updated alignments, only the Australia/VIC01/2020 sequence showed a unique mismatch to the GPS™ probe. Therefore, a full calibration was run using synthetic RNA-genomes from Australia/VIC01/2020 isolate and the resulting Ct values correlated with this obtained from Wuhan-Hu-1 synthetic RNA-genome (Figure 3), indicating that mismatch in the probe is tolerated.

The *in silico* analysis for exclusivity was more complex, showing a wide range of discriminative power for the methods subjected to analysis (Table 1). For instance, the two RT-qPCR designs IP2 and IP4 developed by Institut Pasteur seems to discriminate well between SARS-CoV-2 and other respiratory virus as confirmed for a panel of specimens [14]. The CDC from Atlanta (USA) designed 3 different primer/probes sets named N1, N2 and N3 [15]. We found a low exclusivity in the N3 primer/probe, but a few weeks ago, this set was removed from the panel (https://www.who.int/emergencies/diseases/novel-coronavirus-2019/technical-guidance/laboratory-guidance). Both N1 and N2 showed a good level of mismatching with most coronaviruses except for some Pangolin-CoV sequences which showed very few nucleotide differences. The RT-qPCR proposed by Corman et al., 2020, designed to detect SARS-CoV-2, SARS-CoV and Bat SARS-like-CoV [12], is probably the most used worldwide. They suugested the use of E gene assay as the first-line screening tool, followed by confirmatory testing with the two probes P1 and P2 in the RdRp gene assay. While P1 probe should react with both SARS-CoV-2 and SARS-CoV, P2 probe was considered specific for SARS-CoV-2. Although our *in silico* results confirmed that purpose for P1 (Table 1), the RdRp_P2 assay may also react with some other coronaviruses. The CDC in China developed two RT-qPCR assays for ORF1ab and N genes [16]. Both showed a good overall mismatching to consider they are exclusive, except for some Pangolin-CoV sequences. A similar conclusion may be taken for the N-gene RT-qPCR at the Ministry of Public Health of Thailand [18]. Data of Table 1 indicated that primer/probe of Chu et al., 2020 [17], may be reactive with SARS-CoV-2, SARS-CoV and Bat SARS-like-CoV. The exclusivity of the RT-qPCR design developed by Shirato et al., 2020 [19] clearly resided in the reverse primer as showed 7 mismatches with all SARS related coronaviruses. Finally, Chan et al., 2020 [13] developed three RT-qPCR assays targeting RdRp/Hel, S and N genes of SARS-CoV-2. They selected the RdRp/Hel assay as considered to give the best amplification performance and was tested in parallel with the RdRp-P2 from Charité-Berlin [12]. All positive patients with the RdRp-P2 assay were positive with the RdRp/Hel design. However, 42 patients negative for the RdRp-P2 assay were positive with RdRp/Hel and they found that only RdRp-P2 assay, but not RdRp/Hel, cross-reacted with SARS-CoV culture lysates [13]. Above findings agreed with expected exclusivity derived from the present study. Additional comparative *in vitro* analysis [31] have indicated that primer/probes of ORF1ab from the CDC-China [16] seems the most sensitive, the N2 and N3 assays from the CDC-Atlanta were the most recommended [31]. This partially disagrees our findings as the N3 design may react with other coronaviruses than SARS-CoV-2 (recently removed for the CDC panel). In the study by Arun et al., 2020 [32], the specificity of methods from Charité-Berlin and CDC-Atlanta were tested finding no false positive results but differences in the sensitivity. The most sensitive were N2 (CDC-Atlanta) and E (Charité-Berlin). However, the present study indicates the RT-qPCR for E target may react with different SARS coronavirus. Finally, the kit GPS™ COVID-19 dtec-RT-qPCR Test have shown the highest number of mismatches (i.e., 19-48) for all CoV sequences described so far, including these of Pangolin-CoV which showed a range of 19-31 mismatches. In addition, considerable indels were discerned which enlarge even more the exclusivity of this design.

The GPS™ kit passed the analytical and diagnostic validation according to criteria of the UNE/EN ISO/IEC 17025:2005 and ISO/IEC 15189 (Table 2). The analysis standard curve was repeated a minimum of ten times and average value for all parameters were optimum according to standard limits. For reliability, the coefficient of variation (CV) obtained in all cases for both, repeatability and reproducibility, was always much lower than 10%. The LOD was tested with the usual protocol for 10 copies repeated 15 times with a positive result in all cases (100%). LOQ assays were performed in two sets of 15 tests for both 10 copies of standard templates. The LOQ measurement in both cases was validated with a t-Student test with a confidence interval of 95%. The kit received diagnostic validation at two different reference laboratories (ISCIII, Madrid; and PHE, London). The results shown in Table 3 indicated 100% of diagnostic sensitivity and 100% of diagnostic specificity was assigned. Currently, the kit is being used in several Spanish hospitals and diagnostic laboratories.

Obviously, at the time of designing the RT-qPCR published [12–19], a lack of SARS-CoV-2 genomes available may explain the relatively scarce exclusivity found in some cases. Despite the greater or lesser *in silico* specificity of these primers and probes, due to host specificity of Bat-CoV, Bat SARS-like, Pangolin-CoV, together with the fact of that no human-SARS have been reported since 2004, all positive results obtained would be considered as SARS-CoV-2 infections [17, 33]. However, RNA viruses may exhibit substantial genetic variability. Although efforts were made to design RT-qPCR assays in conserved regions of the viral genomes, variability resulting in mismatches between the primers and probes and the target sequences can result in diminished assay performance and possible false negative results. Primers and probes should be reviewed and updated according to new data, which will increase exponentially during the next few weeks/months.

## ACKNOWLEDGEMENT AND FUNDING

In memory of all the health professionals who have given their lives to care for their patients during this pandemic. We are most grateful to the scientists from the Instituto de Salud Carlos III, Madrid, Spain and the Public Health England, London, UK. This research received no specific grant from any funding agency in the public, commercial, or not-for-profit sectors.

